# Robust differentiation in a synthetic stem-cell circuit

**DOI:** 10.1101/2023.05.01.538905

**Authors:** David S. Glass, Anat Bren, Elizabeth Vaisbourd, Avi Mayo, Uri Alon

**Affiliations:** Department of Molecular Cell Biology, Weizmann Institute of Science, Rehovot, Israel 76100

## Abstract

Differentiation is a process fundamental to multicellularity. In its simplest form, differentiation converts self-renewing stem cells into non-proliferative cells with specified function. This process is inherently susceptible to mutant takeover — mutant stem cells that never differentiate produce excess proliferative daughter cells, driving cancer-like expansion and decreasing the availability of differentiated cells to the organism. It has been proposed that coupling differentiation to an essential trait can select against these mutants by producing a biphasic fitness curve. This would provide mutant stem cells that do not differentiate with a selective disadvantage. However, this theory has yet to be tested experimentally. Here we use “fitness landscape engineering” to design and construct a synthetic biological model of stem cell differentiation in *Escherichia coli* with biphasic fitness. We find that this circuit is robust to mutations as predicted. Surprisingly, its optimal differentiation rate is robust to a wide range of environmental pressures. This environmental robustness is driven by transit-amplifying cells that differentiate and proliferate irrespective of environment. These results provide new interpretations for natural differentiation mechanisms and suggest strategies for engineering robust, complex multicellular consortia.

## Introduction

Differentiation is a fundamental multicellular trait ^1–9^. In adult mammals, many tissues are maintained by ongoing differentiation from stem cells, including the gut, skin, and blood ^10^. Within a tissue, stem cells are a small subpopulation that self-renews indefinitely^2^. The majority of their cell divisions go towards production of differentiated cells, in many cases by first producing transit-amplifying (progenitor) cells, which proliferate rapidly but transiently, ultimately differentiating terminally into the non-dividing functional cells of the tissue ^10–12^. The overall balance of differentiation and self-renewal rates can maintain a constant stem : progenitor : differentiated cell ratio ^2,13–16^.

The level of complexity of differentiation lineages varies broadly from system to system, containing for example a variable number of progenitor stages. A complex example is hematopoiesis, which produces the various blood cell types in a multi-step, branching lineage with multiple types of intermediate progenitor cells ^2,3,14,17^. At the other extreme are simple differentiation processes as in the cyanobacterium *Anabaena* PCC 7120, which forms multi-cellular assemblies but contains only two cell types: self-renewing vegetative cells that fix carbon, and non-dividing heterocysts that fix nitrogen ^18^. When nitrogen is scarce, vegetative cells differentiate directly into heterocysts, which provide nitrogen for the rest of the colony. Such simple differentiation is widespread in bacteria ^19^.

Despite its crucial importance in multicellularity, differentiation is inherently susceptible to loss of function. A mutant stem cell which does not differentiate has an advantage over other stem cells: it reproduces faster because more of its progeny are self-renewing ^20–22^. Such a mutant stem cell will thus outcompete properly differentiating stem cells (Fig. 1A) — a process known as “mutant takeover” ^20^. At a larger scale, this is detrimental to the overall fitness of the multicellular organism due to a concomitant loss of appropriate cell type ratios. For example, in certain leukemias, mutant hematopoietic stem cells with abnormally low differentiation rate give rise to an excess of stem cell descendants ^23,24^. These outcompete non-mutant stem cells in the bone marrow and simultaneously cause a dearth of functional circulating blood cells. Mutant takeover is also seen in synthetic differentiation systems ^25^.

**Figure 1.**
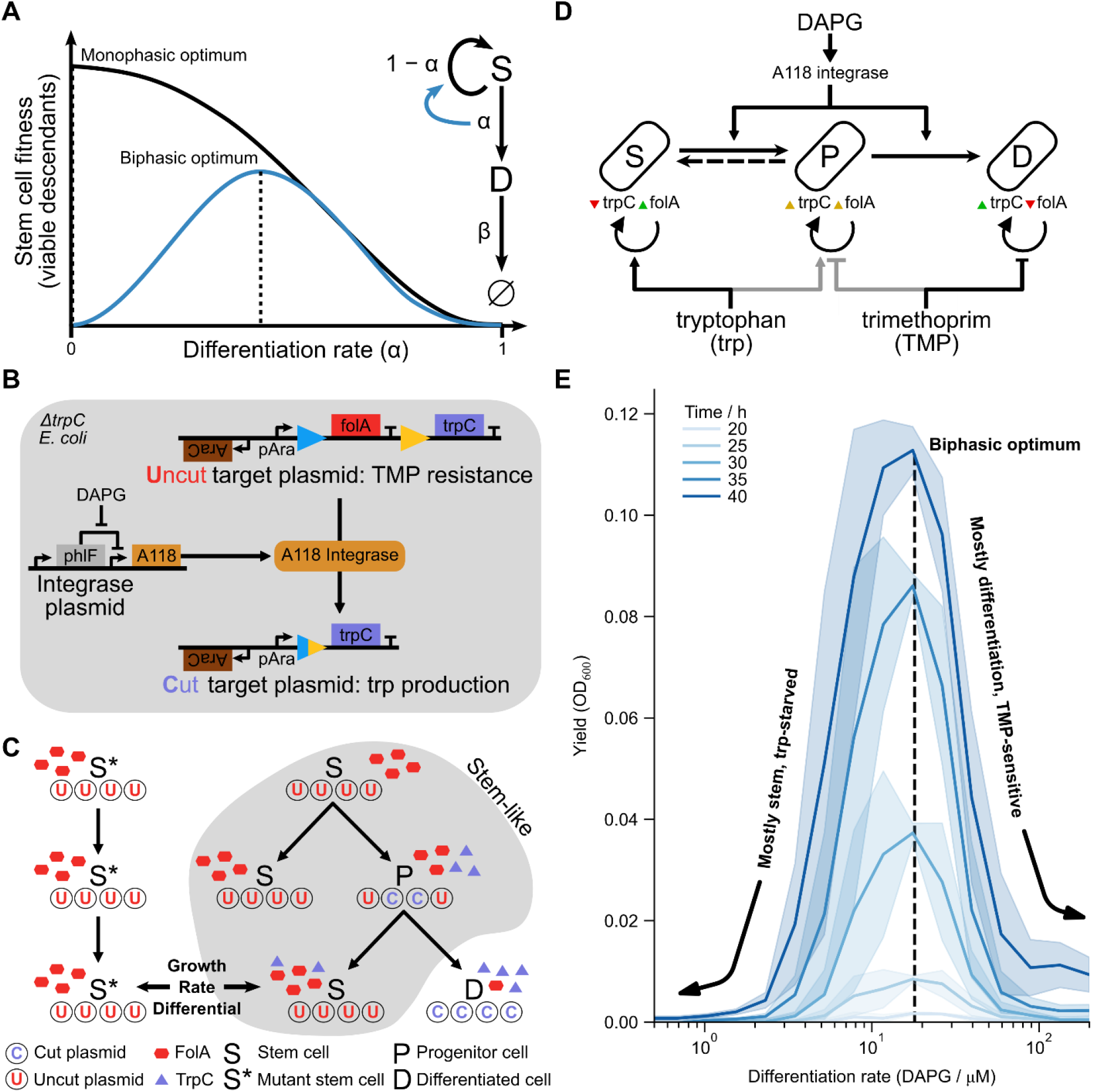
Synthetic differentiating *E. coli* with biphasic control yields a fitness curve with non-zero optimal differentiation rate. **A**. Schematic difference between biphasic and monophasic differentiating circuits. In the monophasic case, zero differentiation yields the maximum fitness. In the biphasic case, a non-zero differentiation rate α yields optimum fitness. **B**. Molecular detail of the biphasic differentiation circuit. Integrase, expressed in all cells, irreversibly cuts a target plasmid, removing trimethoprim resistance gene *folA* and simultaneously inducing expression of the essential tryptophan-producing *trpC*. **C**. Population-level design of the differentiation circuit. Due to random replication and segregation of plasmids, progenitors can give rise to genetically “pure” stem-cell descendants, which nonetheless inherit tryptophan and *trpC* in their cytoplasm. Such recently differentiated stem cells have a fitness advantage over non-differentiating, “mutant” stem cells S*, which have no tryptophan. **D**. The circuit design allows three primary experimental knobs controlled by external factors DAPG, trp, and TMP. DAPG induces integrase expression that differentiates S to P to D, with some reversibility of P to S (see C). Environmental pressures on differentiation can be tuned via external trp and TMP. **E**. The engineered strain indeed shows a biphasic fitness curve as a function of differentiation rate, as induced via DAPG, in low-trp (1.56 μM), high-TMP (25 ng/μL) media. The location of the fitness curve appears fixed over time. Shaded areas are standard deviations across three replicates on separate days.

Given this inherent susceptibility to mutants, there must be control mechanisms in place that prevent loss of stem-cell differentiation throughout adult life in most individuals ^21^. One proposed mechanism is antagonistic pleiotropy ^26,27^: a trait (such as differentiation) which is naively disadvantageous to a cell can be stabilized by having a second essential or beneficial function. In such a case, loss of the ability to differentiate would then give the mutant cell a selective disadvantage, thereby preventing mutant takeover.

Karin, et al. ^20^ expanded the binary concept of antagonistic pleiotropy into a continuous one termed biphasic fitness. In biphasic fitness, a quantitative trait such as differentiation rate increases fitness (i.e., overall proliferative capability) of the stem cell up to a point before the disadvantage takes over and fitness drops. Plotting stem cell fitness versus differentiation rate then includes both a rising phase and a falling phase (hence “biphasic”), with an optimal, non-zero differentiation rate. This is in contrast to the situation in which differentiation only reduces stem cell fitness, resulting in a monotonic dependence of fitness on differentiation rate (Fig. 1A). Biphasic control has been studied in oncogenic mutants ^28–31,32,33^ and population control circuits with no differentiation ^34,35^. In the realm of stem cell differentiation, the concept of biphasic control has yet to be tested experimentally.

One approach to study biphasic control of differentiation is to search for such control within naturally occurring tissues. However, finding such a system can be challenging, as natural biphasic regulation can be convoluted. For example, biphasic regulation can select for differentiation at early times but against differentiation at later times ^27^. This makes identifying natural biphasic regulation far from trivial. In contrast, a synthetic “build-to-understand” approach ^36,37^ can be useful by allowing one to test and design a differentiation circuit and its mutant resistance properties in a controlled setting. One would expect the synthetic biphasic control to maintain differentiation by preventing the spread of non-differentiating stem-cell mutants.

Here we engineer a synthetic stem cell differentiation process with intermediate progenitor states into *Escherichia coli*. We provided this strain with biphasic control, coupling differentiation to expression of an essential gene. The engineered strain provides a controlled biological model to study biphasic stem-cell differentiation. We use competition and evolution experiments to demonstrate the extent of mutant resistance provided by this control.

Furthermore, by growing cultures in a variety of growth conditions, we probe the effect of environmental pressures on growth and differentiation rates. We find that biphasic control provides robustness to mutant takeover. Surprisingly, it is also robust to environmental changes, maintaining nearly the same optimal differentiation rate across a wide range of conditions. This environmental robustness is driven by transient-amplifying states. Thus, we demonstrate that biphasic differentiation can ensure ratios of cell types that are protected both from environmental variation and from mutant takeover.

## Results

### Biphasic circuit based on integrase-driven differentiation

Inspired by natural differentiation, we sought to design a synthetic circuit that irreversibly generates non-growing daughter cells (Fig. 1B-D). The design uses an integrase to cut out a segment of DNA to generate differentiated cells — defined by the absence of this DNA segment. The excised DNA carries antibiotic resistance so that differentiated cells cannot divide in the presence of antibiotic. Biphasic fitness is introduced by engineering the cutting process to induce expression of an essential metabolic gene. Thus, losing differentiation is coupled to loss of this essential metabolic function.

Specifically, we introduced into *E. coli* a large serine integrase, A118, via a plasmid which we call the integrase plasmid. The integrase recognizes a pair of integration sites on a separate target plasmid and irreversibly removes the intervening sequence. We designed this sequence to contain a trimethoprim (TMP) resistance gene, *Pseudomonadota folA*. Differentiation corresponds to the action of the integrase. We define stem cells as those cells containing *folA* on uncut target plasmids and differentiated cells as those lacking *folA*. Thus, stem cells can grow in the presence of TMP whereas differentiated cells cannot.

To achieve biphasic control of differentiation, we engineered the cells so that the act of differentiation, namely cutting out of the DNA segment, induces expression of a gene that provides a fitness advantage. This is done by working in a *ΔtrpC* strain auxotrophic for tryptophan (trp). We designed the target plasmid so that the integrase splices a copy of *trpC* in front of the *folA* promoter, thus coupling differentiation to production of essential tryptophan.

For this design to work, we anticipated that it would be important for the target plasmid to be present in multiple copies (Fig. 1C). With a single copy, stem cells would lack *trpC* and would not grow for lack of tryptophan, whereas the differentiated cells would lack *folA* and would not grow for lack of TMP resistance. We therefore used a medium-copy (p15A) target plasmid.

Likewise, we used a low-copy (pSC101) integrase plasmid to lower the integrase-to-target ratio, aiming to generate a dynamic range of cut vs. uncut plasmids as a function of integrase expression. This departs from previous synthetic uses of integrase for generation of logic functions, which used single-copy targets with high integrase expression to avoid aberrant intermediate states ^38–40^.

With a range of states (i.e., number of cut plasmids) available to cells, stemness now becomes a matter of degree rather than a binary feature — similar to sequential differentiation pathways with multiple progenitors ^2,41^. We thus define a pure stem cell as one with no cut plasmids, a progenitor cell as a cell with an intermediate number of cut plasmids, and a fully differentiated cell as a cell in which all plasmids are cut.

The biphasic design makes use of these multiple states by coupling recent integrase activity to the fitness of each state (Fig. 1C). The integrase produces decreasingly stem-like states by cutting target plasmids. Random replication and segregation of target plasmids into daughter cells can either increase or decrease stemness by producing cells with more or fewer uncut plasmids, respectively. This results in a pool of stem and progenitor cells that are to some extent interchangeable, echoing state transitions in metazoan differentiation lineages ^42,43^.

Pure differentiated cells cannot return to this pool due to the irreversibility of the cutting process. These cells are thus terminally differentiated, and any remnant FolA protein inherited from progenitor parents will be diluted by cell division until growth is stopped by TMP. On the other hand, pure stem cells (lacking cut plasmids) do descend at some rate from progenitors. These stem cells inherit TrpC cytoplasmically, which is then diluted out until further integrase activity induces *trpC* expression. The cytoplasmically inherited TrpC thus provides a fitness advantage to stem cells whose recent ancestors differentiated, as compared to mutant stem cells that have not differentiated for many generations due to loss of integrase function.

The resulting synthetic strain, denoted biphasically differentiating *E. coli* (BDEC), is designed to have several experimental knobs for tuning parameters (Fig. 1D). Differentiation rate is controlled via integrase expression, under control of the inducer 2,4-diacetylphloroglucinol (DAPG), which releases repression by the transcription factor PhlF. Tryptophan and trimethoprim in the media modulate the selection pressure against stem cells and differentiated cells, respectively (Fig. S1). Finally, arabinose controls the overall expression of *folA* and *trpC*.

To test whether the fitness of this strain is indeed biphasic, we grew BDEC in a medium with low trp and high TMP (Fig. 1E). We started with a population of pure stem cells (> 97% uncut plasmids, see Methods) and quantified fitness as the culture yield (measured by optical density) as a function of DAPG concentration and growth time. Optical density was maximal at intermediate integrase induction (∼20 μM DAPG), irrespective of the duration of the experiment, despite a monotonically decreasing population of uncut plasmids (Fig. S2). Without induction, no growth was observed within 48 h. Likewise, a pure differentiated population (pre-cut target plasmid) failed to grow within 48 h. Co-culture of *folA*-expressing cells with *trpC*-expressing cells also did not grow, ruling out cross-feeding of tryptophan or resistance between stem and differentiated cells (Fig. S3).

We conclude that BDEC fails to grow at low integrase induction due to lack of tryptophan, and it fails to grow at high induction due to a lack of TMP resistance. At intermediate integrase induction, we obtain a mixture of cells with varied degrees of differentiation, as evidenced by the presence of both cut and uncut plasmids in the resulting culture (Fig. S2).

Thus, BDEC constitutes a synthetic bacterial analog of a biphasic differentiation circuit. The stem cells can self-renew and generate progenitor cells, which irreversibly generate terminally differentiated cells that do not grow in low trp, high TMP conditions.

### Biphasic differentiation circuit evolves to a well-defined differentiation rate

In the initial experiments described above, the differentiation rate was set externally by the inducer DAPG, which determines integrase expression. Under these conditions, there is an optimal differentiation rate defined by intermediate induction that maximizes yield.

We next asked whether natural selection would stabilize a particular differentiation rate, and whether this rate would be close to the optimal rate that we found by adjusting DAPG. For this purpose, we performed a competition experiment between 11 strains with an identical circuit, but with different levels of integrase achieved by a different A118 ribosome binding site (RBS) in each strain (Fig. 2A). These RBSes, selected from a library of 4096 expected to cover ∼5 orders of magnitude in translation rate (Methods, Fig. S4), were chosen to exhibit a range of integrase expression (Fig. 2B).

**Figure 2.**
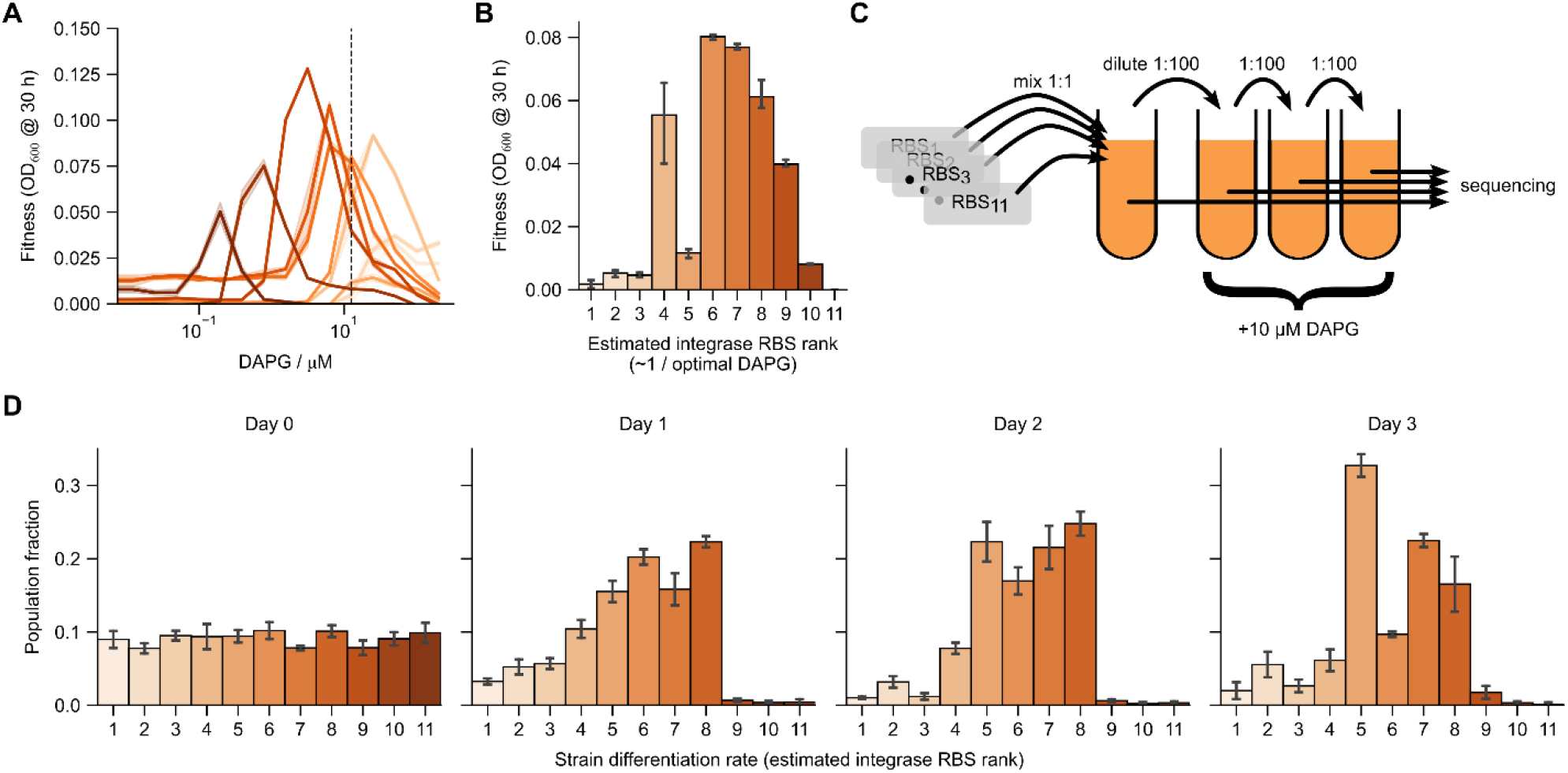
Competition of biphasic strains with different differentiation rates selects for an intermediate differentiation rate close to the optimal rate. **A**. Eleven individual strains with different RBSes of varying strengths each have their own biphasic fitness curve as a function of DAPG, with shifted peaks showing optimization for different externally supplied DAPG. Shaded bands are standard deviations over 3 technical (same-day) repeats. Hue is consistent in subsequent panels to simplify comparison. **B**. The fitness of different strains at a single DAPG concentration (dashed line in A, 12.5 μM) appears approximately biphasic when strains are ordered by rank of integrase RBS rate (differentiation rate), estimated as 1/peak DAPG for each strain. This suggests that competition at a single DAPG concentration should select for intermediate-rate strains. Note that the estimated RBS rates for strains ranked 4 and 5 differ by only ∼40%. **C**. A competition across strains was run by mixing equal concentrations of strains, diluting once per day, and sequencing the RBS region. **D**. Competing the 11 strains by serial dilution over several days shows selection for the strain with optimal, intermediate-level differentiation rate under 10 μM integrase induction. Error bars are standard deviations across 4 independent histories.

Upon induction by DAPG, these strains provided different fitness curves each with a different optimal induction level (Fig 2A). At 12.5 μM DAPG, the yield as a function of ranked RBS strength in fact resembles the biphasic fitness curve of the original strain for a range of DAPG (Fig. 2B), suggesting a fitness advantage to strains with an intermediate differentiation (integrase expression) rate.

To compete the strains, we mixed the 11 strains equally, and grew them in 10 μM DAPG, low-trp, high-TMP medium over the course of several days, diluting the culture 1:100 each day (Fig. 2C, Methods). By sequencing of the evolved cultures, we found that variants of intermediate strength were strongly selected for within 3 days (Fig 2D). The strain population fractions by day 3 ranked closely according to the distance of their optimal DAPG induction from 10 μM (Fig. S5). In particular, the most prevalent strains have peaks closest to 10 μM DAPG.

We conclude that the biphasic synthetic differentiation circuit can evolve to a defined differentiation rate close to the optimum. The circuit provides a selective disadvantage to variants that increase or decrease integrase expression. In particular, it provides a selective disadvantage to variants with very low integrase activity, which can be considered mutant stem cells that do not differentiate.

### Differentiation rate is robust to environmental pressures

The experiments described so far were performed in a single environment (medium), with low trp and high TMP (see Methods). We next asked how changing the environment affects the fitness curve as a function of differentiation rate (DAPG concentration), and in particular how it affects the optimal differentiation rate. This is a robustness question, because changes in the environment alter the selective pressure on the two arms of the biphasic curve.

We first repeated the induction curve experiment in three additional extreme environments in which we added minimal or saturating amounts of trp and TMP (–trp –TMP, +trp +TMP, +trp –TMP = ± 25 μM trp ± 1.56 ng/μL TMP) (Fig. 3A). Without trp or TMP (–trp – TMP), the fitness curve rose monotonically with DAPG (Fig. 3A, solid blue line), reflecting the fact that increasing differentiation provides increased trp production without the cost of reduced TMP resistance. With abundant trp and TMP (+trp +TMP), the fitness curve decreased approximately monotonically (Fig. 3A, dashed orange line) — increased differentiation only causes loss of resistance, the abundant trp removing any selective pressure to induce *trpC*. With abundant trp and no TMP (+trp –TMP), the fitness function was approximately flat (Fig. 3A, solid orange line), reflecting no selective pressure on production of either trp or TMP resistance.

**Figure 3.**
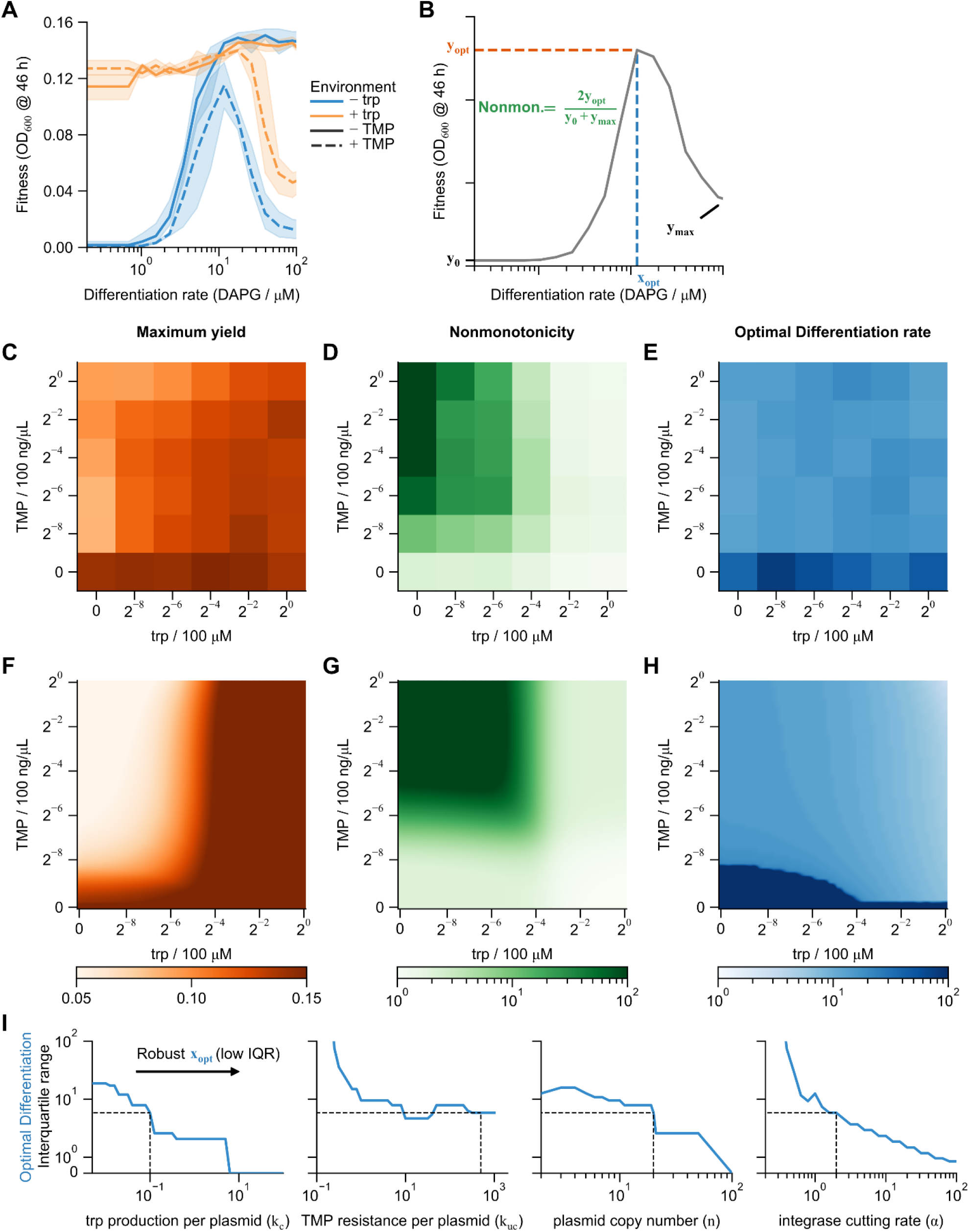
Biphasic differentiation is robust to environmental conditions. **A**. Fitness curves with saturating levels of trp and/or TMP show expected monotonically increasing, monotonically decreasing, biphasic, and flat trends. **B**. For a broad range of environmental conditions (trp and TMP), fitness curves were quantified by the peak fitness and corresponding optimal differentiation rate (in units of DAPG), as well as a measure of nonmonotonicity of the fitness curve. **C-E**. Heat maps of the three metrics show graded responses in yield and monotonicity, but nearly constant optimal differentiation rate except in the complete absence of TMP. **F-H**. An interchangeable cell-type logistic growth model (Methods) qualitatively reproduces the response of these three metrics. **I**. Interquartile range of optimal differentiation rate (*x*_*opt*_) across all trp/TMP environments as a function of model parameters. Robust *x*_*opt*_ (low IQR, indicating little environmental influence) occurs with intermediate to high plasmid copy number and *folA* and *trpC* production per plasmid comparable to sourcing from the environment. Together these imply a need for many progenitor states whose growth rate is independent of environment.Cutting rate also needs to be faster than growth rate (α > 1). Dashed lines reference the parameter values used in F-H. Values in A,C-E are averages of repeats on 3 separate days. Bands in A are standard deviations over those repeats

To understand how such divergent fitness curves interconvert in less extreme environments, we repeated the induction curve experiment in 36 environmental combinations of 6 levels of trp and 6 levels of TMP, each ranging in concentration across four orders of magnitude. For each environment, we quantified the maximum yield (*y*_*opt*_), optimal differentiation rate (*x*_*opt*_), and nonmonotonicity of the fitness curve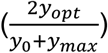 (Fig. 3B). As expected, optimum yield increased with increasing trp and decreasing TMP (Fig. 3C), reflecting the decreasing selective pressures. Likewise, nonmonotonicity increased with decreasing trp and increasing TMP (Fig. 3D). In most intermediate conditions, the fitness curve showed a peak at intermediate DAPG concentration (Fig. 3E).

Notably, the optimal differentiation rate varied by only ∼1.7-fold across environmental conditions (trp/TMP), except in complete absence of TMP. This suggests that the optimal differentiation rate in the biphasic system is robust to environmental conditions that change over four orders of magnitude.

To understand the origin of this robustness we developed a mathematical model that describes the growing culture as logistic growth of interconverting cell types (see Methods for further details). Using a set of parameters estimated from data and literature (Table S1), the model agrees with the data qualitatively (Fig. 3F-H). In particular, it recapitulates the robustness of optimal differentiation rate, as well as the trends in maximum yield and nonmonotonicity.

To test which parameters controlled this robustness, we re-ran the simulations with different parameter values (Fig. 3I, S6) and quantified the variation in optimal differentiation rate by the interquartile range of *x*_*opt*_ across all environments. We found that the optimal differentiation rate is robust as long as plasmid copy number is sufficiently high (≳10), *trpC* and *folA* expression per plasmid are strong enough to be comparable to sourcing trp or removing TMP from the environment, the cutting rate is faster than division, and “leaky” growth rates are low. As these conditions have only a lower or upper bound, none require fine tuning. Thus, the biphasic circuit provides robust differentiation rate as a function of environment, in a manner robust to the circuit parameters.

The mathematical requirements for environmental robustness of differentiation rate can be interpreted as a need for many progenitor states. These intermediate states are transient but evade selective pressures (having both abundant tryptophan production and TMP resistance in our system). This suggests that a large number of progenitor cells, which occupy intermediate differentiation states, provide an environmentally robust differentiation rate in biphasic differentiation cascades by temporarily ignoring environmental pressures.

### Differentiation is resistant to mutant takeover in long-term evolution experiments

We next asked whether the biphasic circuit is robust to mutations that change the differentiation rate. For example, complete loss of differentiation would be analogous to cancer-like mutations that fail to differentiate. To do so, we grew the BDEC strain (Fig. 1E) in –trp +TMP (10 ng/μL TMP) media with moderate integrase induction (8 μM DAPG), diluting 1:100 into fresh media approximately once per day (Fig. 4A, Methods). Assaying the fraction of cut plasmids by qPCR as a proxy for differentiation rate, we found no significant change in differentiation within 24 transfers, or ∼160 generations (Fig. 4B). The cultures indeed appeared to approach a reproducible steady-state differentiation fraction. This indicates that through selective pressure, biphasic control maintains differentiation rate.

**Figure 4.**
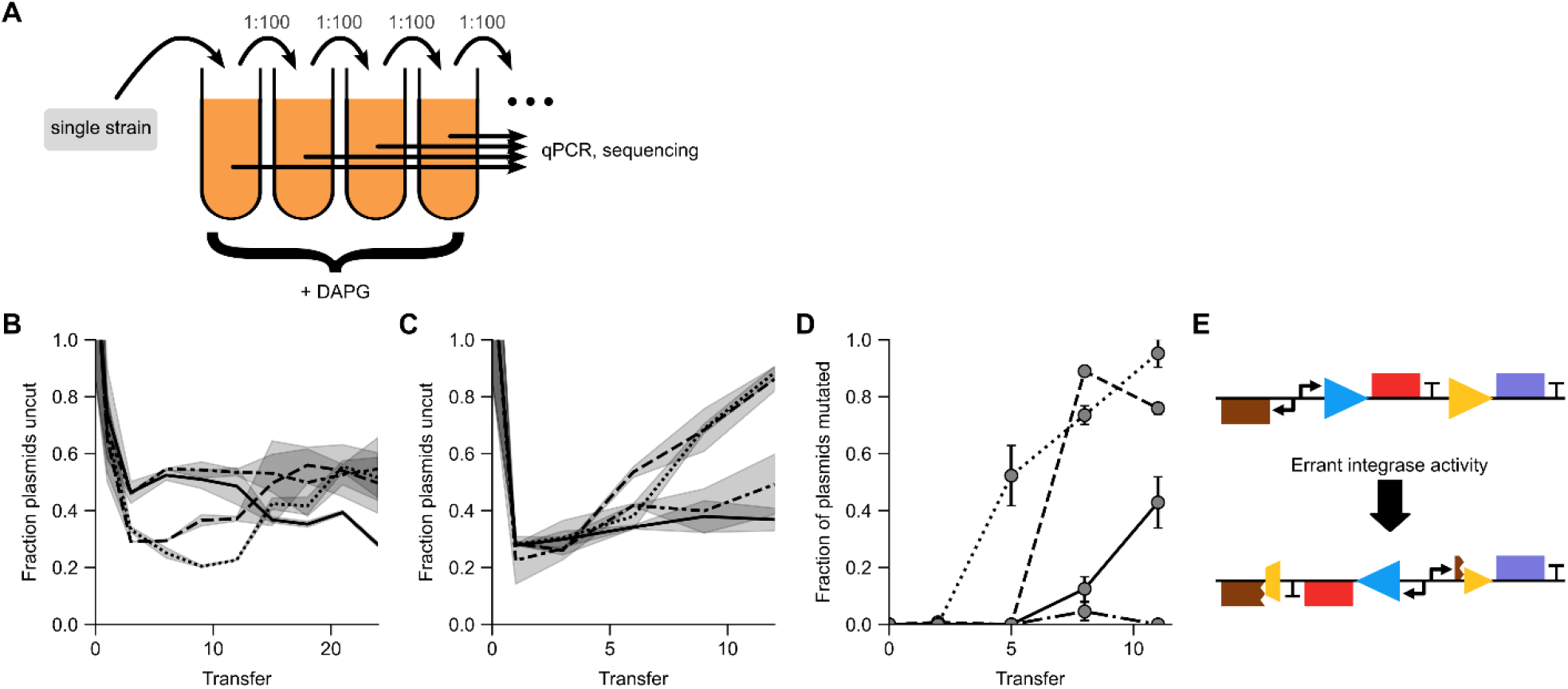
Biphasic differentiation is maintained in long-term evolution but lost under weak selective pressure due to decoupling of the biphasic control. **A**. Schematic of evolution experiment, starting with the original BDEC strain cultured in low-pressure media (10 ng/μL TMP), passaged approximately once per day, and quantified by qPCR and nanopore sequencing. **B**. Fraction of uncut plasmids (approximate number of stem cells) in an evolved culture does not change significantly over 4 weeks (∼160 generations) with moderate integrase induction (8 μM DAPG), appearing to reach a reproducible steady-state differentiation fraction.**C**. Under high integrase induction (20 μM DAPG), the fraction of uncut plasmids approaches 1 within ∼1.5 weeks (∼45 generations), indicating a non-differentiating mutant with selective advantage. **D**. Sequencing of evolved cultures from C indicates a mutant increasing in abundance over the ∼1.5 weeks of the high-induction experiment in 3 out of 4 independent replicates. **E**. Detailed analysis of the mutant sequence shows that the mutant simultaneously expresses both trp production (*trpC,purple*) and TMP resistance (*folA, red*) via errant integrase action, which inverted the *folA* and pAra promoters, while breaking production of AraC. It is presumed that *folA* expression relies on the native (genomic) copy of AraC. Bands in B,C are standard deviations of two technical repeats. Error bars in D are bootstrapped from sequence counts (Methods). Line styles in B-D refer to independent histories. Colors in E match Fig. 1B.

We then increased the selective pressure on differentiation by repeating the experiment with higher integrase induction (20 μM DAPG). In this case, we found nearly complete loss of differentiation for 2 out of 4 repeats within 7 transfers, or ∼45 generations (Fig. 4C), suggesting mutant takeover by non-differentiating clones. To determine what mutations had occurred, we sequenced the culture plasmids, assuming that relevant mutations were most likely to occur within the circuit construct itself. Indeed, we found that both strains contained inversion mutations that broke the biphasic coupling between loss of *folA* expression and induction of *trpC*, allowing their simultaneous expression (Fig. 4D). Similar mutations in fact were found increasing but still at low frequency in the two repeats without complete loss of differentiation, suggesting that this sidestepping of biphasic control via decoupling is reproducible. The inversion matches up at one end with the integrase target site, indicating that this mutation was mediated by errant integrase activity, which can invert DNA targets with appropriately oriented target sites ^40^.

## Discussion

We used a “build-to-understand” approach to explore the extent to which biphasic control can protect stem cell differentiation against evolutionary and environmental pressures. We built a synthetic differentiation system in *E. coli*, in which an integrase irreversibly removes antibiotic resistance in stem-like cells (differentiation) while simultaneously inducing production of an essential amino acid (biphasic control). This biphasic control strain selected for a specific differentiation rate in competition and provided protection against mutant takeover in long-term evolution experiments. The mutant resistance was lost in conditions that strongly drove differentiation. Unexpectedly, the biphasic mechanism also provided robust cell type ratios in the face of variations in growth conditions. By comparison with a mathematical model, we showed that this environmental robustness is due to many transit-amplifying (progenitor) states that are not modulated by exogenous factors. Thus, biphasic control resists takeover by mutants that lose differentiation, and maintains ratios of stem and differentiated cells across conditions.

### Comparison to previous synthetic differentiation

Mutant resistance in our system was lost when driving differentiation at a high rate, corresponding to a large loss of stem cells. At lower induction of differentiation, the ratio of stem to differentiated cells was maintained over ∼160 doublings. A recent system using synthetic differentiation in *E. coli* without biphasic control similarly showed loss of differentiation at high differentiation rates ^25^. While that system sustained differentiation over ∼90 doublings at low differentiation rates, further comparison is difficult due to the differences between the two systems in design and quantification. Interestingly, both systems failed at high induction with mutations caused by errant recombinase behavior, its DNA-editing capability apparently increasing the effective mutation rate.

### Comparison of synthetic E. coli differentiation to natural stem-based tissues

From our modeling work, we found that transient-amplifying (progenitor) stages are crucial to an environmentally robust differentiation rate. Such differentiation cascades are common in mammalian tissues ^2,3^. In such tissues, as in our synthetic model, progenitors function as amplifiers, which reproduce faster than either stem cells or terminally differentiated cells. For systems that need to alter differentiation rate in response to the environment, one would expect few intermediate states. This matches, for example, the cyanobacterium Anabaena PCC 7120, which contains no progenitors and differentiates only when nitrogen is scarce ^18^.

The conditions in our system that lead to loss of differentiation are likewise suggestive for analogies to mammalian diseases. With mutational loss of differentiation, the stem-differentiated cell type ratio increases progressively. This is similar to the increasing overabundance of stem cells and decreasing number of differentiated cells seen before the onset of idiopathic pulmonary fibrosis ^44,45^, osteoarthritis ^46,47^, and leukemia ^14,23,24^. Changing environmental conditions in our system that favor stem cells (increased tryptophan) could lead to similar non-genetic effects with potentially more abrupt loss of differentiated cells once the fitness of pure stem cells exceeds the biphasic benefit gained from differentiation. This may be akin to niche-induced cancer, in which local environmental changes modify the stem cell balance between differentiation and self-renewal ^48^.

### Fitness landscape engineering

The present work focuses on specifying fitness as a function of quantitative genetic and environmental factors, a process we term “fitness landscape engineering.” This differs from standard synthetic biology efforts to decouple circuit behavior from a cellular “chassis” ^49^. In fact, if one considers the biphasic differentiation circuit as a logic circuit, it encodes a simple function: replacing *folA* and *trpC* in our circuit with green and red fluorescent proteins would produce a simple BUFFER/NOT function. It is in connecting both outputs of this logic circuit to cellular fitness that we obtain a fitness landscape biphasic with respect to a quantitative genetic trait, differentiation rate. Assessing this landscape is more complicated than assessing logic behavior, and requires growing cells both short- and long-term in a variety of environmental conditions. In this work, this workflow enabled us to produce a robust differentiation circuit and discover that transit-amplifying states enable this robustness.

We hypothesize that a similar same design-build-test-learn workflow could be applied, *inter alia*, to improve multicellular engineering ^37,50,51^, biocontainment ^52–54^, population control ^34,55,56^, modulation of antibiotic tolerance ^57,58^, cancer treatment ^59,60^, and evolutionary stability of synthetic circuits ^61^ or biomolecule production ^62,63^.

The ultimate failure mode of our biphasic control circuit suggests directions for improving fitness landscape engineering. In particular, the mutants that took over at high integrase induction were unexpected, but perhaps predictable in retrospect. This suggests that in future engineering efforts, potential mutations might be computationally predicted and designed around. Complete exploration of all mutants in a fitness landscape is infeasible computationally and experimentally. However, algorithms that combine automated feature annotation with *in silico* evolution could significantly improve the ability to build mutant-resistant circuits ^64–66^. In the case of the mutants found here, such an algorithm would minimally need to (1) generate inversions, (2) predict transcription/translation rate of rearranged sequences, and (3) provide a model of growth under given environmental conditions.

### Future work

In future work, it would be interesting to study the spatiotemporal behavior of the biphasic differentiation circuit. Natural stem cell-based tissues often have defined structures. For example, stem cells are evenly spaced among differentiated cells in the fly gut, despite drastic changes in the size of the tissue during development and starvation ^13^. In mammals, stem cells are localized in a colonic crypt ^41^. How stem cell lineages, fitness landscapes, and spatial location interact to maintain such structure could be potentially assayed using our synthetic model if grown in a structured environment.

One could also introduce population control^3,55,67^ into the synthetic differentiation system. In contrast to the biphasic differentiation control proposed in Karin, et al.^20^, the biphasic control in this work is cell-autonomous and decoupled from population feedback control. This feedback could be introduced via induction of differentiation in stem cells by differentiated cells. The cell-autonomous nature of our biphasic control could help in preventing cheaters that fail to secrete or sense diffusible feedback signals ^55,68^.

More generally, it would be interesting to study other proposed population and mutant control mechanisms by engineering them into *E. coli*. Mechanisms like reciprocity ^69^, spring-and-ceiling ^70^, or autoimmune surveillance of hyper-secreting mutants (ASHM) ^71^ have precise theoretical predictions that could be assayed in controlled biological contexts via fitness landscape engineering. We thus predict that, in addition to engineering evolutionarily stable multicellular consortia, fitness landscape engineering will have wide-ranging applications in understanding natural mechanisms for controlling tissue growth in multicellular organisms.

## Acknowledgments

The authors would like to thank H. Kim, T. Milo, V. Jayaraman, Y. Yang, M. Raz, and the Alon lab for helpful feedback and comments on the manuscript. Funding was provided by European Research Council (ERC) under the European Union’s Horizon 2020 research and innovation program (grant agreement No. 856587). D.S.G. was funded as a member of the Zuckerman Postdoctoral Scholars Program. U.A is the incumbent of the Abisch-Frenkel Professional Chair.

## Methods

### Strains and plasmid construction

To prepare a *ΔtrpC::KanR* strain, the knockout region was transferred to MG1655 (CGSC #6300) from Keio collection strain JW1254 (BW25113 *ΔtrpC770::KanR*) using P1 phage transduction ^72^. Plasmids were constructed by Genscript, and in some cases gel-purified to avoid plasmid multimers — particularly in the target plasmid — and verified by Plasmidsaurus. Initial sequences not produced via synthesis were sourced from Addgene (Roquet, et al.^40^) or by PCR from the MG1655 genome (*trpC*). To avoid A118 cutting target plasmids during strain construction, target plasmids were always transformed into strains already containing the integrase plasmid with its associated repressor, as described in Roquet, et al. To verify that no target plasmids were cut during cloning, the transformed stock was sequenced and analyzed by nanopore sequencing as described below. A full list of strains used in this work is provided in Table S2.

### Growth conditions

Standard growth conditions were designed to provide a tryptophan-free baseline. Cultures were grown overnight in LB, washed 5x in M9 minimal medium (42 mM Na_2_HPO_4_, 22 mM KH_2_PO_4_, 8.5 mM NaCl, 18.5 mM NH_4_Cl, 2 mM MgSO_4_, 0.1 mM CaCl) by centrifugation at 6000 rpm in a tabletop centrifuge for 3 min and replacing the supernatant with fresh M9 each time. Starting cultures were then diluted back ∼1:500 into (unless otherwise stated) M9 + 0.4% arabinose supplemented with 50 μg/ml spectinomycin and/or 50 μg/ml ampicillin (for strains containing integrase and/or target plasmids, respectively) and supplemented with the specified concentrations of inducer (DAPG) and “environmental conditions” (trp, and TMP). For short-term growth experiments, final growth cultures were prepared in 96-well assay plates by a Tecan FreedomEvo liquid handler in a final volume of 200 μl and covered with 50 μl mineral oil. Plates were then grown in incubators at 37°C shaken at 6 Hz. Every ∼30 min the plates were transferred by robotic arm to an Infinite M200Pro plate reader, which recorded the OD_600_. Reported OD values are background subtracted per-well against the mean of the first few timepoints. Stocks of trp, TMP, and DAPG were stored at -20°C. Long-term stocks of trp were stored at –80°C.

### Library selection

An initial integrase-plasmid library (pDSG589L) was constructed by Genscript with a randomized 6-bp region within the RBS of A118, predicted by the RBS calculator ^73^ to range in translation rates over ∼5 orders of magnitude. Electrocompetent cells of the *ΔtrpC* strain were prepared by serial washing in cold water (chilled 40 mL exponential-phase cells on ice 15 min, centrifuged at 1000 g for 10 min, removed supernatant and transferred to 1.5 mL conical tube, washed 2x at 1000 g for 5 min each, resuspended in 100 μl water + 10% glycerol). The pDSG589L library was then transformed by electroporation using Bio-Rad MicroPulser with default Ec2 settings. Around 10^4^ – 10^5^ colonies were collected, grown for ∼2 h in LB and stored with 25% glycerol at –80°C. Before storing as glycerol stocks, a sample was prepared directly as electrocompetent cells and transformed with the target plasmid (pDSG545). A total of 576 colonies were picked and grown overnight in 1 mL LB in deep 96-well plates, washed, and grown as described above. Twelve strains were selected as described in Fig. S4. Individual integrase plasmids in each strain were collected by miniprep (Qiagen). The RBS region of each was sequenced by Sanger sequencing, revealing one “strain” (thereafter removed from future experiments) to be a mixture of two strains. The remaining 11 minipreps were transformed by heat shock into the *ΔtrpC* strain and tested by selection to ensure only integrase plasmids had been transformed. The target plasmid (pDSG545) was then subsequently transformed into each of the 11 strains.

### Competition

The 11 selected strains were grown overnight in LB, washed in M9 as described above, OD-normalized, and mixed in equal volumes. A sample of the mixture was stored as glycerol stock. The mixed culture was then diluted 1:100 into four 1 mL-cultures in a deep 96-well plate with M9 + arabinose + ampicillin + spectinomycin media as above supplemented with 0.25 μM trp and 500 ng/μL TMP and 10 μM DAPG. Cultures were grown overnight covered in breathable film. A sample of overnight culture was stored as a glycerol stock. Cultures were then diluted 1:100 into fresh media on subsequent days without any additional trp. DAPG was added fresh each day to avoid degradation. Glycerol stocks were PCRed in a ∼1 kb region surrounding the RBS and sequenced using nanopore sequencing (Plasmidsaurus).

### Evolution

For the evolution experiment, an overnight culture of BDEC was grown and washed as described above. The washed culture was diluted 1:100 into four 1mL-cultures in M9 + arabinose + ampicillin + spectinomycin media as above supplemented with 0.25 μM trp and 10 ng/μL TMP and 8 μM or 20 μM DAPG. Each day, glycerol stocks were made from overnight cultures. Cultures were then diluted 1:100 into fresh media without trp. Media was stored for the duration of the experiments in 4°C without DAPG, which was added fresh each day. Once per week cultures were transferred every 2 days instead of every day.

### Nanopore sequencing and analysis

Nanopore sequencing was performed by Plasmidsaurus. Sequences were aligned to known plasmid references using minimap2 2.21-r1071 and processed using samtools 1.12 and seqkit 2.2.0. Subsequent analysis was performed in python 3.9.7 using Bipython 1.79 and pysam 0.16.0.1. Mutations were called using NanoSV version 1.2.4 and longshot. The mutation frequencies (Fig. 4D) were calculated as the number of sequences supporting the called variant (output by NanoSV) divided by the sequencing depth calculated by samtools at the locus defined of the inversion point (i.e., confidence interval output by NanoSV). Bootstrapped errors were simulated as the standard error from 10000 repeats of drawing, with replacement, *y* mutants from *n* sequences where *y* is the number of called mutants and *n* is the sequencing depth.

### qPCR

Glycerol samples were diluted 1:10 into water and mixed with Applied Biosystems Fast SYBR Green mastermix (#4385612) and 4 pmol each primer. Each sample was probed in duplicate both within *folA* and within *AmpR* (on plasmid backbone). Samples were placed in an Applied Biosystem MicroAmp Fast Optical 96-well reaction plate (#4346906), covered in MicroAmp Optical Adhesive Film (#4311971), and run on an Applied Biosystem QuantStudio 3 qPCR machine. Within each qPCR plate, a sample was run from the glycerol stock before the evolution experiment (Day 0), a sample of a pre-excised strain (lacking *folA*) and a strain with no integrase plasmid (fully uncut). Raw fraction uncut was calculated as 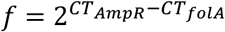 Reported fraction uncut plasmids were normalized for primer efficiency as (*f − f*_*no foLA*_)/*f*_*no integrase*_.

### Model

The growth model is an extension of logistic growth, which is generally modeled as

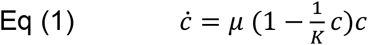

with *c* the concentration of a single species of bacterium, *μ* the growth rate of that bacterium, and *K* the carrying capacity of the culture. In our model, we extend this single-species logistic growth to logistic growth of multiple interconverting cell types. We write this as:

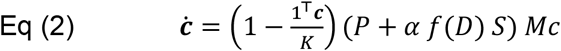

where **c** is a vector of cell numbers *c*_*x*_, indexed by number of cut plasmids *x*. The carrying capacity *K* is assumed to apply to the entire culture, and hence divides 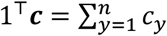 is a diagonal matrix containing the growth rate of each species (*M*_*xx*_ = *μ*(*x*), other entries zero). *P* and *S* are matrices that describe how the various species interconvert — *P* is a stochastic matrix driving random distribution of plasmids to daughter cells during division, and *S* determines the conversion of *c*_*x*_ to *c*_*x*+1_ via differentiation. The parameter α is the maximum integrase cutting rate, and the function 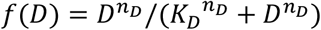 is an increasing Hill function of DAPG concentration *D* with half-maximal concentration *K*_*D*_ and cooperativity *n*_*D*_.

We now describe the matrices *P, S*, and *M* in detail.

#### The replication/distribution matrix P

We model the random replication and segregation of plasmids by the product of two processes: (1) binomial replication of *n* plasmids of which *y* are cut to produce *z* cut plasmids, with probability *y*/*n* of replicating a cut plasmid of in each case; and (2) hypergeometric selection of *n* plasmids of which *x* are cut from the 2*n* of which *y* + *z* are cut. All possible *z* are summed over, giving a probability of producing a cell with *x* cut plasmids from a cell with *y* cut plasmids:

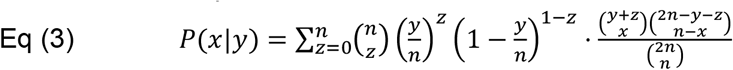

The production of cells with *x* cut plasmids is then 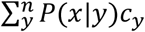 and in matrix form provides a term 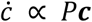 *P****c*** where *P*_*xy*_ = *P*(*x*|*y*).

#### The differentiation matrix S

To model differentiation, we note that differentiation removes *c*_*x*_ and produces *c*_*x*+1_. Thus, the rate of change due to differentiation for *c*_*x*_ is given by 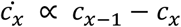 and in matrix form can be written as *S****c***, where *S* is defined as

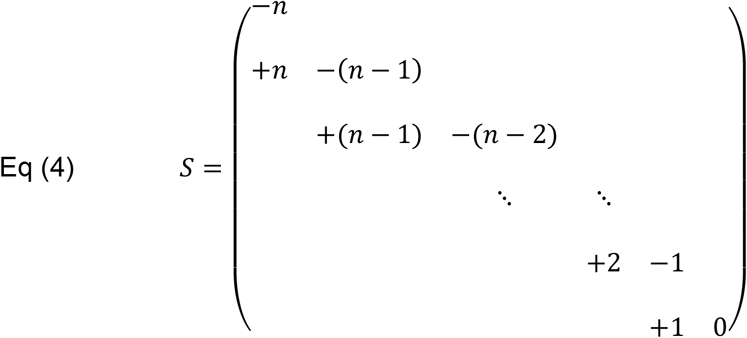

#### The growth-rate matrix M

We approximate the growth rate as a product of Michaelis-Menten functions with leakage. This approximates the logic that trp and cut plasmids (*trpC*) increase growth rate while TMP decreases growth rate with protection provided by uncut plasmids (*folA*):

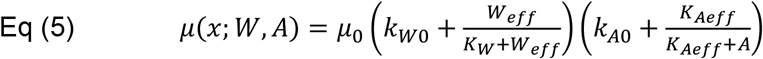

where effective tryptophan *W*_*eff*_ = *W* + *K*_*W*_*k*_*c*_*x* and effective TMP protection *K*_*Aeff*_ = *K*_*A*_(1 + *k*_*uc*_(*n − x*)). The parameters *W* and *A* represent the trp and TMP concentrations, respectively, n the plasmid copy number, *K*_*W*_ and *K*_*A*_ Michaelis-Menten coefficients providing the concentration scales. The parameters *k*_*c*_ and *k*_*uc*_ determine the effectiveness of cut and uncut plasmids, respectively in providing tryptophan and resistance. The growth rate for each species is independent, contributing a term 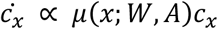, and in matrix form can be written as *Mc* with diagonal matrix *M* :

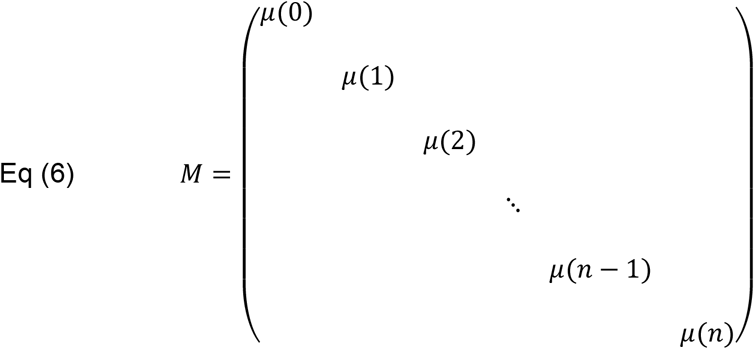

In component form, Eq (2) can be written as:

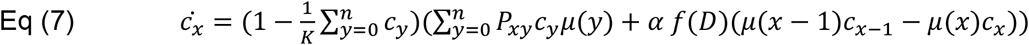

### Simulations

Equations were numerically integrated in Mathematica 12 using NDSolve. For the initial simulations, 62 log-spaced concentrations of TMP and trp were used, with 82 log-spaced concentrations of DAPG for each condition ranging from 0.1 to 1000 (in addition to 0). For the parameter sweeps, the same ranges were used, but with 32 different concentrations of trp and TMP, and 42 concentrations of DAPG. The values of other simulation parameters and how they were estimated are provided in Table S1.

