## Supplementary Information for "Robust differentiation in a synthetic stem-cell circuit"

**Includes:**

**Figures S1 – S6**

**Tables S1 – S2**

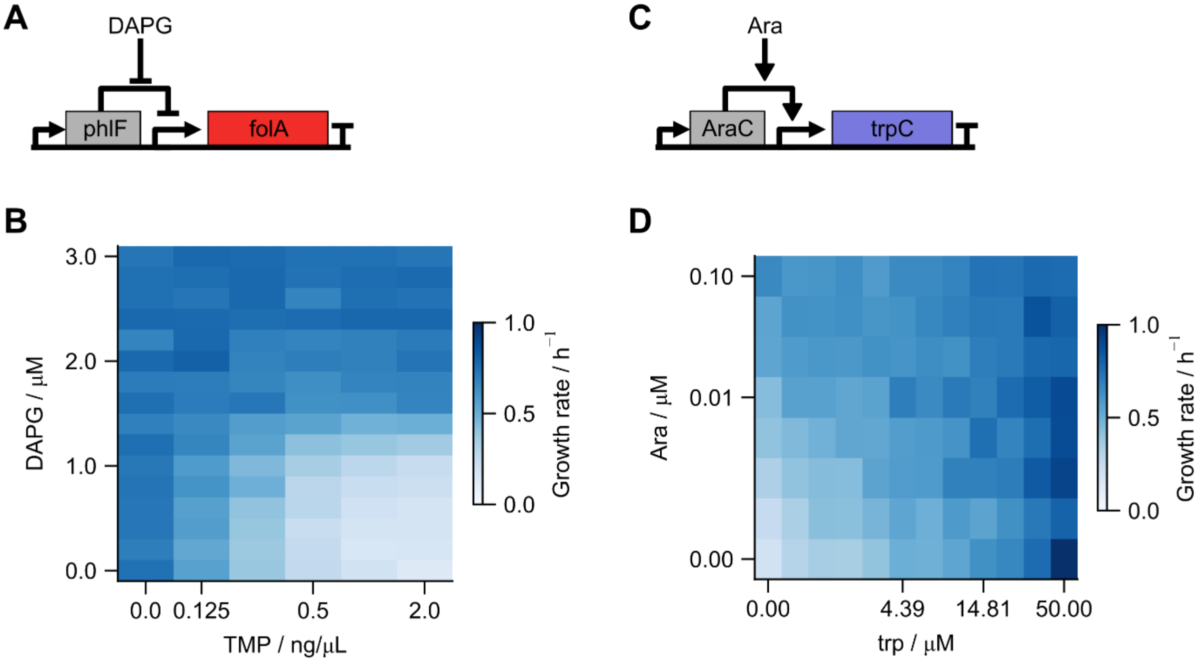


**Figure S1. Direct control of *trpC* and *folA* allow measurement of 2D input functions based on induction of the two genes and their environmental-pressure counterparts.** The scale parameters $K_A$ and $K_W$ for simulation are approximately equal to the concentration that gives half-maximal growth rate at zero induction. Cultures were grown in M9 + 0.4% glucose.

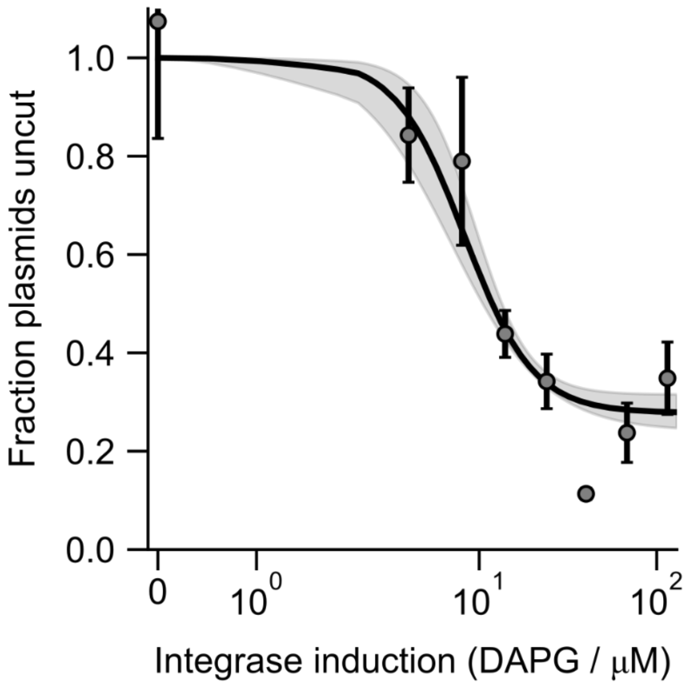


**Figure S2. Fraction of uncut plasmids decreases with increased induction by DAPG in low trp (1.5 μM), high TMP (25 ng/μL) culture.** Note that peak fitness is around 10 – 20 μM DAPG. Curve is a fit of the data (assuming the point at 40 μM DAPG as an outlier) to the Hill function $fraction = k_0 + (1 - k_0)\frac{K^n}{K^n + DAPG^n}$ with $k_0 = 0.28 \pm 0.04$, $K = 8.25 \pm 1.24$, and $n = 2.24 \pm 0.61$. Points are mean ± standard error.

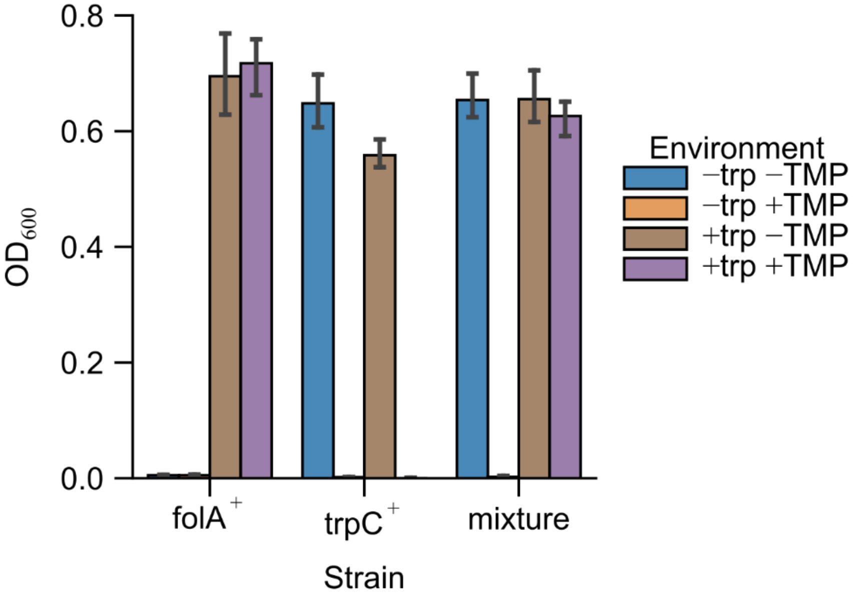


**Figure S3. Cross-feeding between *trpC*-producing and *folA*-producing strains is not significant.** Strains expressing either *folA* or *trpC* under arabinose induction (MG1655 Δ*trpC* pDSG546 or pDSG550) were grown in M9 + 0.4% glucose + 0.4% arabinose + 300 mM cAMP with the specified environment (± 200 μM trp ± 100 ng/μL TMP). Strains grew to a consistent final OD, in permissive environments only. Co-culture of 1:1 mixtures of the strains failed to grow in the environment requiring both gene products (–trp +TMP).

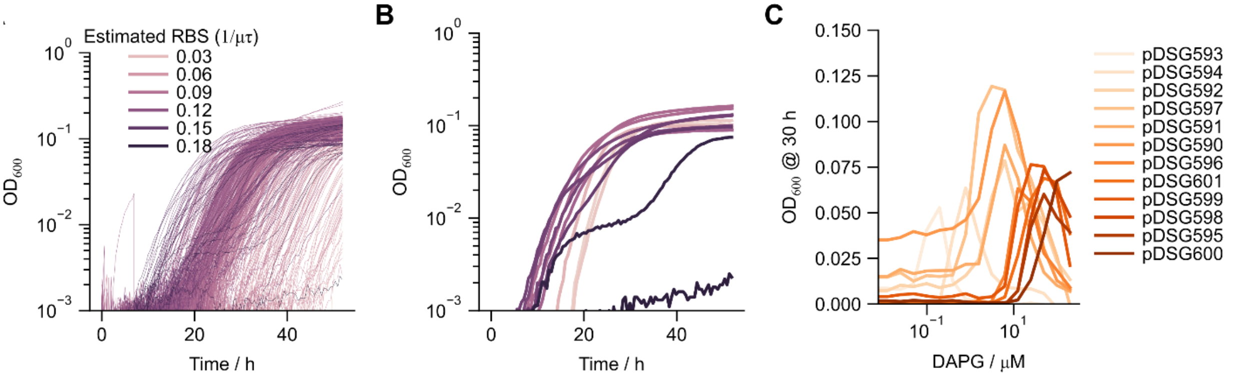


**Figure S4. Overview of the screening process for the 11 strains used in the competition assay.** The strains were selected from 596 colonies of a randomized 6-nt RBS library (maximum 4096). Delayed logistic functions were fit to each strain to provide an estimated RBS rank (~ 1 / (lag * growth rate)) under –trp +TMP growth. Twelve strains with a range of estimated RBSes were selected and grown in a range of DAPG concentrations to confirm shifted fitness peak location. The integrase plasmid of each one of these twelve was sequenced; one of the twelve was determined to be a mixture of two strains and was removed from further experiments.

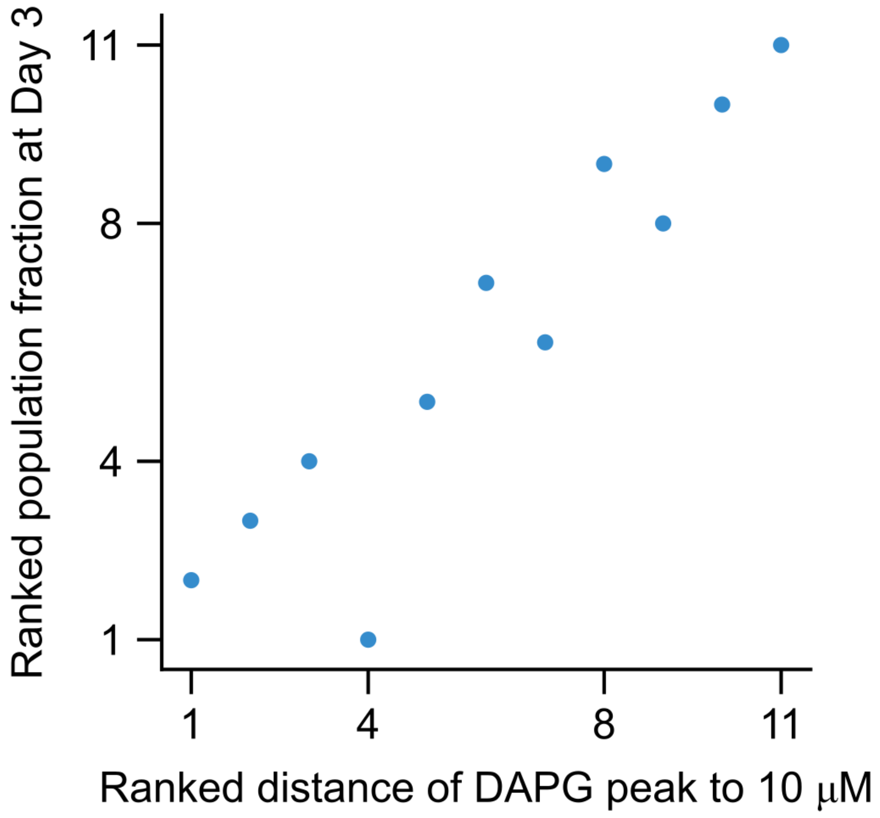


**Figure S5. Population fraction of strains are predictably selected in competition.** The ranked population fraction of each competed strain (Fig. 2E) after competition highly correlates with how close the optimal DAPG concentration of the strain (Fig. 2C) is to the concentration (10 μM) used in the competition assay. Distance is calculated as $|log\ (DAPG_{max}/10)|$. Population fraction rank is defined as lowest rank for highest population fraction.

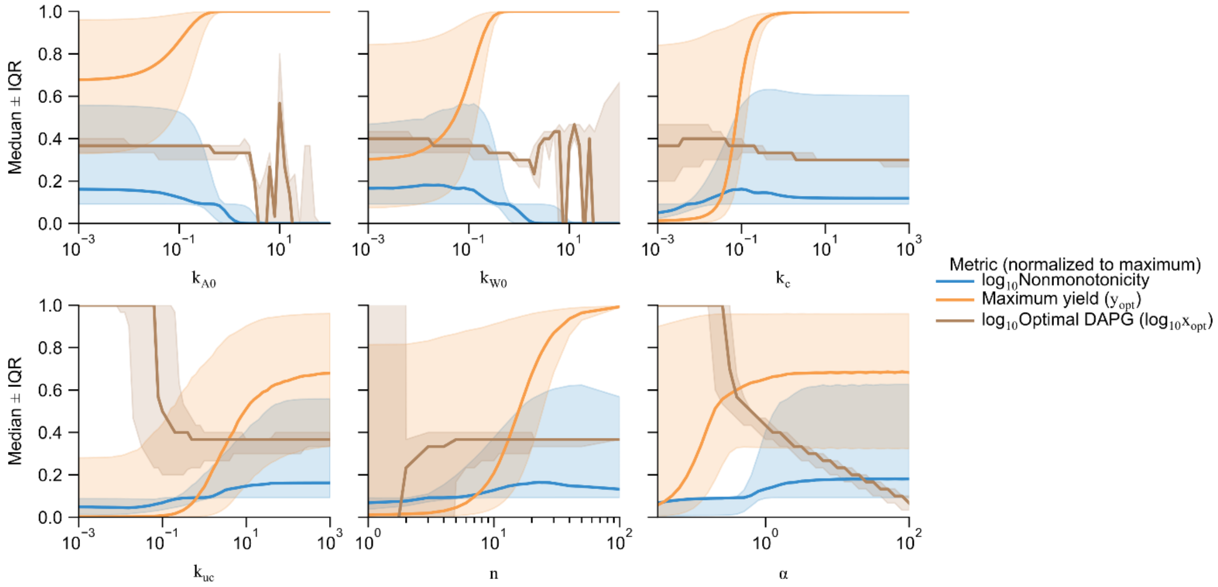


**Figure S6. Extended data on parameter sweeps.** Each metric is normalized to its maximum value across all simulations and then the median and interquartile range (IQR) are calculated. High values of $k_c$, $k_{uc}$, $n$, and alpha give low range (shaded bands) of optimal DAPG while maximum yield and nonmonotonicity are not low (which would indicate a non-growing culture or flat fitness curve, respectively). The same is true for low growth-rate “leakage” $k_{W0}$ and $k_{A0}$.

**Table S1**
Parameter values used for simulation data (Fig. 3F-H)

| Parameter | Value | Description | Estimated from |
|---|---|---|---|
| $k$ | 0.2 | Maximal OD | Fig. 3A |
| $c_0$ | 0.00002 | Initial OD | Fig. 3A, Methods |
| $t_{max}$ | 30 h | Time extent of simulation | Methods |
| $k_D$ | 30 μM | Half-maximal DAPG induction | Fig. S2 |
| $k_W$ | 10 μM | Half-maximal induction of growth by trp | Fig. S1 |
| $k_A$ | 1 ng/μL | Half-maximal reduction of growth by TMP | Fig. S1 |
| $n$ | 20 | Target plasmid copy number | Literature[74] |
| $\mu_0$ | 0.5 / h | Maximal growth rate (in absence of leakage) | Fig. S4 |
| $n_D$ | 2 | Hill coefficient of integrase induction by DAPG | Fig. S2, Literature[75] |
| $\alpha$ | 2 | Integrase cutting rate relative to growth rate | Chosen to match Fig. 3C-E |
| $k_c$ | 0.1 | Production of trp per cut plasmid | Chosen to match Fig. 3C-E |
| $k_{uc}$ | 500 | Production of TMP resistance per uncut plasmid | Chosen to match Fig. 3C-E |
| $k_{W0}$ | 0.1 | Contribution to growth rate with no trp or *trpC* | Chosen to match Fig. 3C-E |
| $k_{A0}$ | 0 | Contribution to growth rate with high TMP and no resistance | Assumed |

**Table S2**
Strains used in this study

| Strain | Description | Used in |
|---|---|---|
| DSG1 | MG1655 *ΔtrpC::KanR* | Background strain |
| DSG1 pDSG578 pDSG545 ("BDEC") | DSG1 integrase plasmid + target plasmid | Figs. 1,3,4,S2 |
| DSG1 pDSG545 | No-integrase control | Fig. 4 |
| JM109 pDSG553 | Pre-cut target | Fig. 4 |
| DSG1 pDSG589L pDSG545 | RBS library | Fig. S4 |
| DSG1 pDSG590-601 pDSG545 | 12 competition strains (pDSG592 mixture left out) | Figs. 2, S5 |
| MG1655 pDSG488 | DAPG-induced *folA* | Fig. S1 |
| DSG1 pDSG467 | Arabinose-induced *trpC* | Fig. S1 |
| DSG1 pDSG546 | Arabinose-induced *folA* | Fig. S3 |
| DSG1 pDSG550 | Arabinose-induced *trpC* | Fig. S3 |
